# Kinetic analysis of strand invasion during *C. elegans* meiosis reveals similar rates of sister- and homolog-directed repair

**DOI:** 10.1101/2025.01.10.632442

**Authors:** Antonia Hamrick, Henry D. Cope, Divya Forbis, Ofer Rog

**Affiliations:** School of Biological Sciences and Center for Cell and Genome Sciences, University of Utah, Salt Lake City, UT 84112

**Author notes:** equally contributing authors (Antonia Hamrick, Henry D. Cope).

## Abstract

Meiotic chromosome segregation requires reciprocal exchanges between the parental chromosomes (homologs). Exchanges are formed via tightly-regulated repair of double-strand DNA breaks (DSBs). However, since repair intermediates are mostly quantified in fixed images, our understanding of the mechanisms that control the progression of repair remains limited. Here, we study meiotic repair kinetics in *Caenorhabditis elegans* by extinguishing new DSBs and following the disappearance of a crucial intermediate - strand invasion mediated by the conserved RecA-family recombinase RAD-51. We find that RAD-51 foci have a half-life of 42-132 minutes for both endogenous and exogenous DSBs. Surprisingly, we find that repair templated by the sister chromatid is not slower than repair templated by the homolog. This suggests that differential kinetics are unlikely to underlie ’homolog bias’: the preferential use of the homolog as a repair template. We also use our kinetic information to revisit the total number of DSBs per nucleus - the ‘substrate’ for the formation of exchanges - and find an average of 40 DSBs in wild-type meiosis and >50 DSBs when homolog pairing is perturbed. Our work opens the door for analysis of the interplay between meiotic repair kinetics and the fidelity of genome inheritance.

**Key points:**

- By extinguishing new meiotic DSBs, we define the lifetime of a key repair intermediate.
- We find similar rates of meiotic DNA repair templated by the sister and the homolog.
- Kinetic information allows calculation of the total number of meiotic DSBs in *C. elegans*.

## Introduction

The segregation of the parental chromosomes (homologs) into the gametes at the end of meiosis is aided by genetic exchanges, or crossovers. Crossovers physically link the homologs, allowing them to correctly align on the metaphase plate during the first meiotic division. Crossovers form by repairing programmed double-strand DNA breaks (DSBs), which are catalyzed by the conserved topoisomerase-related protein Spo11 (SPO-11 in worms; (1, 2)). Importantly, only a subset of DSBs are repaired as crossovers (3, 4), whose formation requires the use of a specific repair pathway and repair template.

Crossovers form by the homologous recombination machinery, which restores the genetic information on the broken DNA by copying it from an intact homologous sequence. Most meiotic DSBs are repaired via homologous recombination. Indeed, when alternative repair pathways, such as non-homologous end-joining, are eliminated there are little to no meiotic phenotypes in worms (5–8). Homologous recombination relies on the interaction between a resected, single-strand DNA adjacent to the DSB and an intact, double-strand template, from which information is copied. Downstream of these so-called strand-invasion intermediates, repair proceeds along several possible pathways, with a tightly regulated subset of intermediates eventually forming crossovers (4).

Both the homolog and the sister chromatid contain homologous sequences with which to engage in strand invasion. While only homolog-directed repair leads to crossover formation, sister-templated repair can take place during meiosis (9–15). Since sister-directed repair does not lead to formation of crossovers, differential regulation of homolog- *versus* sister-directed repair is critical for generating gametes with a complete genome complement. However, the details of such differential regulation remain poorly understood.

During early stages of homologous recombination, the DNA adjacent the break is resected to expose single-stranded DNA that is then coated with RecA-family recombinases (RAD-51 in worms). This protein-DNA filament then acts as a ’tentacle’ that searches for homology and forms strand-invasion intermediates with a double-stranded template (16–19). During meiosis in *Caenorhabditis elegans*, RAD-51 foci mark DSBs from the time they are resected until the formation of downstream repair intermediates (18, 20, 21). However, it is unknown whether different repair scenarios - such as various repair templates - affect this duration. This gap in our knowledge makes it challenging to interpret changes to the number of RAD-51 foci in fixed images, which is one of the most widely applied readouts for meiotic DSB repair. This challenge has also confounded attempts to assess the total number of DSBs in wild-type and perturbed meiosis. Here, we overcome these limitations by controlling the influx of DSBs, allowing us to quantify the lifetime of RAD-51 foci.

## Materials and methods

### Worm strains and growing conditions

Worms were grown as in (22). All analyzed worms were age-matched, adult hermaphrodites (24 hours post-L4). Worms were grown at 20°C, except strains carrying *syp-1^K42E^*, which were maintained at 15°C (permissive temperature), and were grown at 25°C from hatching prior to immunofluorescence. Auxin plates (1mM auxin) were prepared as in (23). All worms used in this work contained the *SPO-11-AID* allele (23) and the *TIR1* ubiquitin ligase driven by the *sun-1* promotor (23). For the SPO-11 shut-off experiments, worms were moved to auxin plates at the beginning of the experiment (time-point 0 hours). For x-ray irradiation experiments, worms were grown on auxin from hatching. The lack of endogenous DSBs was verified by RAD-51 staining (<0.1 RAD-51 foci per nucleus prior to irradiation). Irradiation was performed as previously described using W-(ISOTOPE) source set to 112mV (9). Worms were irradiated on plates for 150 and 300 seconds, for 1000 and 2000 rads, respectively. For analysis of *syp-3* worms, we used homozygous pharyngeal-GFP-negative progeny of balanced heterozygous mothers (ROG459). The following strains were used:

ROG297: *meIs8 [pie-1p*::*GFP*::*cosa-1 + unc-119(+)] II; spo-11(ie59[spo-11::AID::3xFLAG]) ieSi38 [sun-1p::TIR1::mRuby::sun-1 3’UTR + Cbr-unc-119(+)] IV*

ROG212: *meIs8 [pie-1p*::*GFP*::*cosa-1 + unc-119(+)] II; spo-11(ie59[spo-11::AID::3xFLAG]) ieSi38 [sun-1p::TIR1::mRuby::sun-1 3’UTR + Cbr-unc-119(+)] him-8(tm611) IV*

ROG400: *meIs8 [pie-1p*::*GFP*::*cosa-1 + unc-119(+)] II; zim-2(slc4) spo-11(ie59[spo-11::AID::3xFLAG]) ieSi38 [sun-1p::TIR1::mRuby::sun-1 3’UTR + Cbr-unc-119(+)] IV*

ROG459: *syp-3(ok758) I / hT2 [bli-4(e937) let-?(q782) qls48] I;III spo-11(ie59[spo-11::AID::3xFLAG]) ieSi38 [sun-1p::TIR1::mRuby::sun-1 3’UTR + Cbr-unc-119(+)] IV*

ROG497: *spo-11(ie59[spo-11::AID::3xFLAG]) ieSi38 [sun-1p::TIR1::mRuby::sun-1 3’UTR + Cbr-unc-119(+)] IV; syp-1^K42E^ (slc11) V*

ROG498: *mIn1 [mIs14 dpy-10(e128)] / + II; spo-11(ie59[spo-11::AID::3xFLAG]) ieSi38 [sun-1p::TIR1::mRuby::sun-1 3’UTR + Cbr-unc-119(+)] IV*

zim-2 *allele construction*

*zim-2* was disrupted using CRISPR/Cas9 in the SPO-11-AID background (*meIs8 II; spo-11(ie59[spo-11::AID::3xFLAG]) ieSi38 IV*). CRISPR was performed as in (24) using a *dpy-10* co-conversion marker, and guide RNA 5’-UGUUUCACUAAGUACAACUC-3’. The repair template introduces a premature stop codon after the 75^th^ amino acid. The mutation was verified by PCR and Sanger sequencing, and the new allele was designated *slc4*. The allele recapitulated the *zim-2(tm574)* null phenotype (25): reduced fertility, weak Him and 7 DAPI-staining bodies in diakinesis.

### Immunofluorescence

Immunofluorescence was done as in (26), using overnight incubation at 4°C with the following primary antibodies: guinea pig anti-HTP-3 ((1:500; (27)), rabbit anti RAD-51 ((1:10, 000; (28)), goat anti-SYP-1 ((1:300); (28)). The following secondary antibodies (Jackson ImmunoResearch; 1:500 dilution) were incubated for 2 hours at room temperature: Alexa647 donkey anti-rabbit, Cy3 donkey anti-guinea pig; Alexa488 donkey anti-goat. Images were acquired as z-stacks using a 63x 1.40 NA oil objective on a Zeiss LSM 880 AiryScan microscopy system in FastAiry mode for analysis of *him-8* and irradiated worms. *zim-2, syp-3, mIn1/+* and *syp-1^K42E^* were acquired using AiryScan mode. Image processing was conducted using the ZEN (Blue 3.9, Zeiss).

### Image analysis

The gonad was segmented and divided into seven zones using a custom ImageJ script available upon request. RAD-51 intensity was adjusted to minimize signal outside the nuclei. Counts were performed on maximum intensity projections. Unpaired chromosomes (Figs. 4 and S2) were identified based on asynapsed axes (i.e., lack of SYP-1 staining). The number of nuclei rows in each zone are: zone 4 - 8.4; zone 5 – 8.3; zone 6 – 6.9 (N = 7). Maximum intensity projections are shown throughout.

### Statistical analysis

Exponential fits were performed in Prism 9.5 (GraphPad), using the “one phase decay” option, constraining the decay coefficient to be positive and setting the plateau to 0.

## Results

### SPO-11 shut-off reveals RAD-51 foci lifetime

We analyzed the data in (29), which used a functional auxin-regulated SPO-11 (SPO-11-AID; the strain also expresses the plant E3 ligase TIR1 under a germline promotor; (2, 23, 30)) to prevent the formation of new DSBs and then followed the fate of remaining repair intermediates over time (Fig. 1B). The rate at which RAD-51 foci disappeared – due to their progression to downstream repair intermediates – allowed us to quantify their lifetime. The experiment uses the *C. elegans* oogenic gonad, where all stages of meiosis are presented in a linear fashion, and divides it into 7 equal zones, with the majority of DSBs and RAD-51 foci occurring in zones 4 and 5, where nuclei are in early- and mid-pachytene (Fig. 1A; (18, 19)). The rate of nuclei movement in the gonad – about one nuclei-row per hour (31–35) – entails only limited nuclei movement between zones during the 4-hour time course.

**Figure 1:**
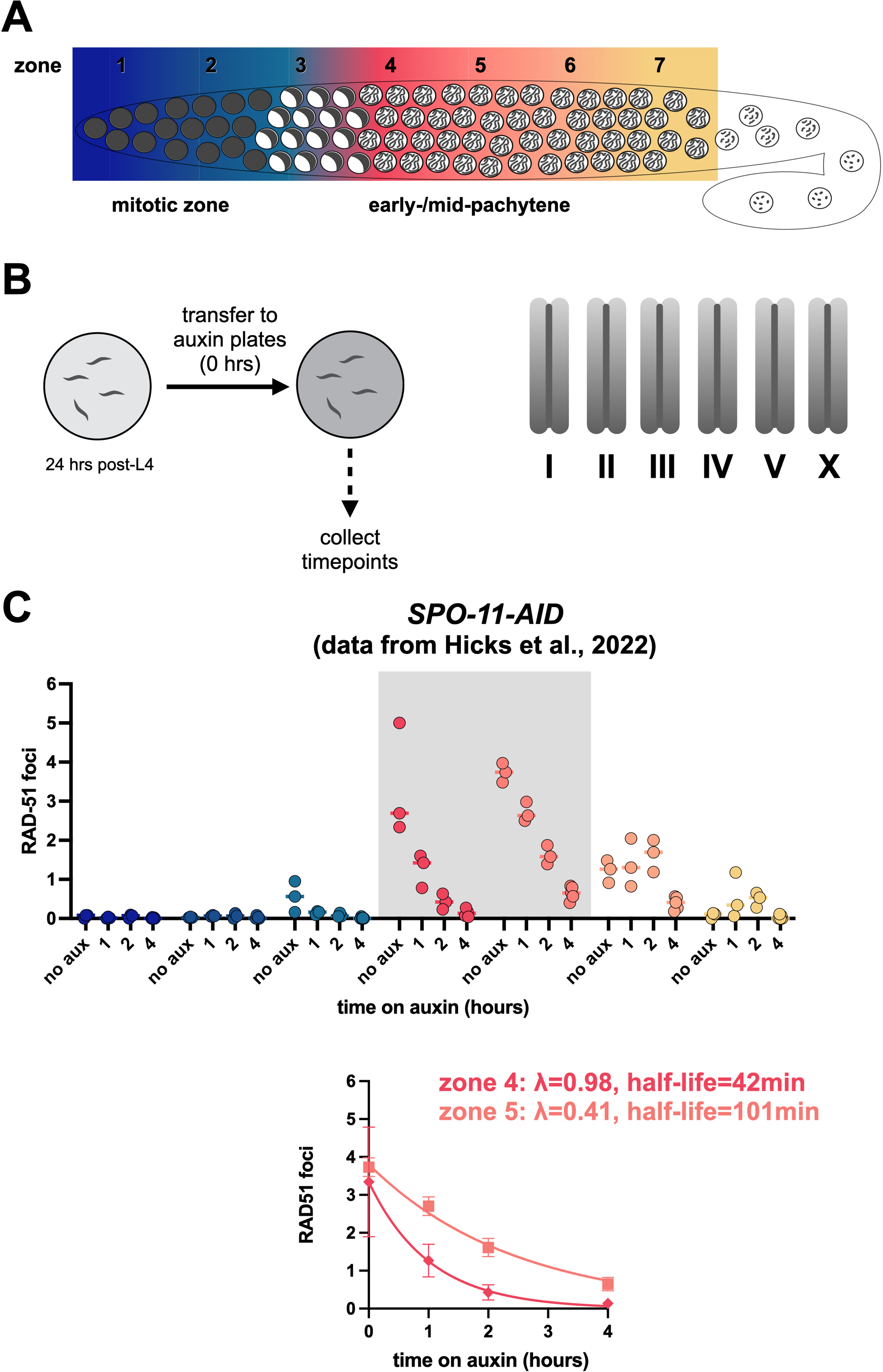
SPO-11 shut-off reveals RAD-51 foci lifetime. (A) Diagram of the *C. elegans* gonad, with meiosis progressing from left to right. Entry to meiosis occurs in zone 3. (B) Diagram of the SPO-11 shut-off experiment. At timepoint 0 hr, adult hermaphrodites (24 hr post-L4) are moved to auxin plates. Right, diagram of the *C. elegans* hermaphrodite karyotype, which includes 5 pairs of autosomes (labelled I - V) and a pair of X chromosomes. (C) Left, RAD-51 foci number per nucleus in *SPO-11-AID* animals following 0, 1, 2 and 4 hours on auxin, colored by gonad zones, as shown in panel A. Data is from (29). Gray shading indicate zones where decay rate was calculated. Bottom, exponential decay of RAD-51 foci number in zones 4 and 5. The half-life was calculated as *ln*(*2*)*/λ*. Error bars indicate standard deviation.

The precise time when the influx of RAD-51 intermediates is halted in this experiment depends on two factors. First, the point when DSB formation is halted as a consequence of SPO-11 degradation on auxin, the rate of which is variable for different targets (23). Second, the time it takes to process new DSBs to allow RAD-51 loading, including removal of the SPO-11 protein adduct and resection to expose single-stranded DNA (4). While the exact duration of these intervals is unknown, the drastic reduction in foci number within one hour of placing SPO-11-AID worms on auxin (Fig. 1C) suggests that these processes together are completed in less than an hour. This duration is consistent with the analysis of meiotic recombination in yeast using physical assays (about an hour separates the appearance of DSBs and single-end invasions, an intermediate downstream of strand-invasion; (36, 37)). Nonetheless, this delay suggests that our calculations below, which assume these processes occur instantaneously, are overestimating RAD-51 foci lifetime.

We modeled the disappearance of RAD-51 foci as a first-order decay process, where *C* is the number of RAD-51 intermediates, *t* is the time in hours, and *λ* is the exponential decay constant:

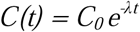

By fitting exponential decay curves, we found that the half-life of RAD-51 foci in control worms is 42 and 101 mins for zones 4 and 5, respectively (*λ* = 0.98, 0.44). Throughout, we limited our analysis to zones with >2 RAD-51 foci per nucleus, since the small starting number of intermediates in other zones prevented robust analysis.

### Similar repair kinetics of radiation- and SPO-11-induced DSBs

We also analyzed strand-invasion kinetics by using an orthogonal approach to control DSB influx. We subjected worms lacking endogenous DSBs (*SPO-11-AID* worms grown on auxin from hatching; hereafter *spo-11(-)*) to x-ray irradiation and monitored RAD-51 foci number over time (Fig. 2A). Radiation-induced DSBs are structurally distinct from SPO-11 DSBs in that they do not necessarily involve protein-DNA adducts (38). However, this difference is unlikely to affect the analysis we carry out here, which monitors post-resection events. Importantly, radiation-induced DSBs are repaired using similar machinery as endogenous DSBs, and recapitulate many of its properties: repair of radiation-induced DSBs allows the formation of crossovers (2), and these crossovers are regulated, much like crossovers generated by the repair of SPO-11-induced DSBs, to number exactly one per homolog pair (3, 24).

**Figure 2:**
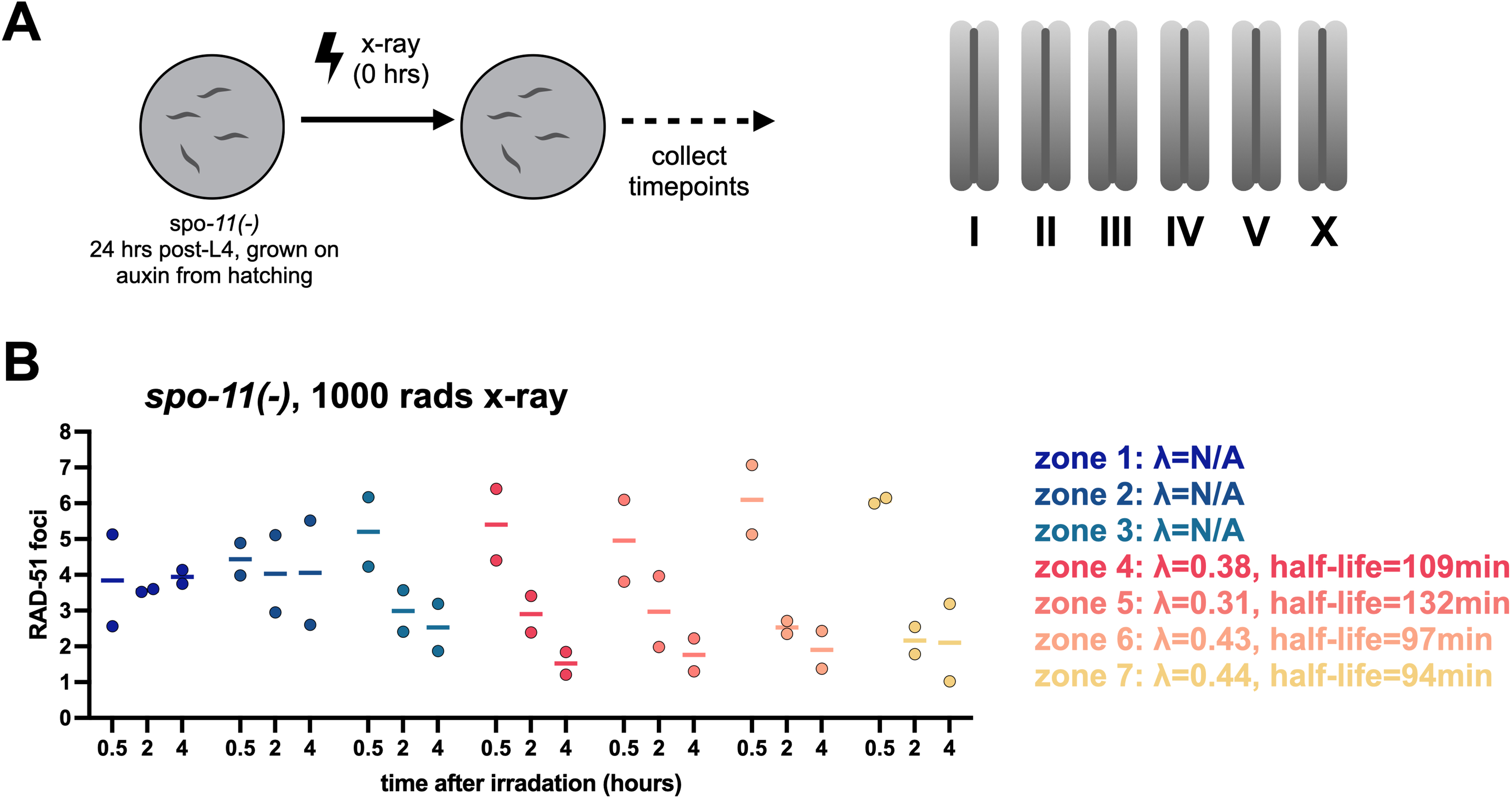
RAD-51 foci lifetime is similar for endogenous and x-ray-induced DSBs. (A) Diagram of the irradiation experiment. 24 hr post-L4 *spo-11(-)* hermaphrodites (SPO-11-AID worms grown on auxin from hatching) were irradiated at timepoint 0 hr. Worms were dissected at the indicated timepoints after irradiation. Right, diagram of the *C. elegans* hermaphrodite karyotype. (B) RAD-51 foci number per nucleus in *spo-11(-)* animals 0.5, 2 and 4 hours after irradiation, colored by gonad zone as in Fig. 1.

The RAD-51 foci half-lives for x-ray-induced DSBs range from 94-132 mins in zones 4-7 (Fig. 2B) indicating that DSBs from irradiation follow similar repair kinetics as endogenous DSBs. Unlike SPO-11-induced DSBs, whose location is tightly regulated (39, 40) x-ray-induced DSBs form at random locations and are thought to be spread evenly across the worm genome. The overall similarity in RAD-51 foci lifetimes between both kinds of DSBs suggests that genomic location has a limited effect on strand-invasion kinetics. The similarity also suggests that the kinetics of DSB formation – occurring over a 2.5-5 minutes-long irradiation or over many hours for SPO-11-induced DSBs (29) – has a limited effect on strand-invasion kinetics. Similarly, the similar repair kinetics throughout most of meiotic prophase (zones 4-7 in the gonad), suggests that meiotic progression has limited effect on strand-invasion kinetics.

A notable difference between repair kinetics of SPO-11- and radiation-induced meiotic DSBs is the number of remaining RAD-51 foci at the end of our 4-hour time course (e.g., 1.5 foci 4 hours after irradiation of *spo-11(-)* worms *versus* 0.2 foci after 4 hours on auxin in SPO-11-AID animals, zone 4). This result suggests that the repair of a small subset of DSBs is differentially regulated, due to their structure or genomic location.

The irradiation experiments allowed us to analyze repair irrespective of meiotic progression. Interestingly, we observed much longer RAD-51 lifetimes in zones 1 and 2, which represent a proliferative stem-cell-like compartment where nuclei undergo mitotic divisions (zone 3, which includes both mitotic and meiotic nuclei also exhibited slow repair; see also images in (21)). This difference might be explained by the meiotic regulation of the homologous recombination machinery, which is geared towards the successful formation of crossovers.

### RAD-51 foci lifetime during sister-directed repair

We wished to test whether the template used during DSB repair influences the lifetime of strand-invasion intermediates. In the conditions tested above, DSBs undergo both sister- and homolog-directed repair, limiting our ability to distinguish between the two. Importantly, meiotic DSBs form throughout the genome regardless of homolog accessibility, and these DSBs are eventually repaired by homologous recombination, as evident by the intact chromosomes at the end of meiosis (9, 13, 19). We therefore used meiotic perturbations where some or all chromosomes fail to partner with their homolog to examine the repair of DSBs that can only engage in sister-directed repair.

We analyzed two conditions that prevent inter-homolog interactions throughout the genome. During meiosis, the homologs align (’synapse’) through the assembly of a conserved chromosomal interface called the synaptonemal complex (4, 41–43). We analyzed *syp-3* worms (*syp-3(ok758*) *SPO-11-AID*) which completely lack a synaptonemal complex, resulting in splayed homologs that are paired only at a small region near their ends (Fig. 3A; (7, 19, 43)). As previously reported, *syp-3* worms exhibit a higher overall number of RAD-51 foci compared to controls, and these foci persist until later stages of meiosis (Figs. 3B and S1; (7)). However, RAD-51 foci numbers decreased rapidly with time on auxin. The decay rates in zones 4, 5, and 6 were 46, 143, and 163 minutes, respectively, similar to control worms.

**Figure 3:**
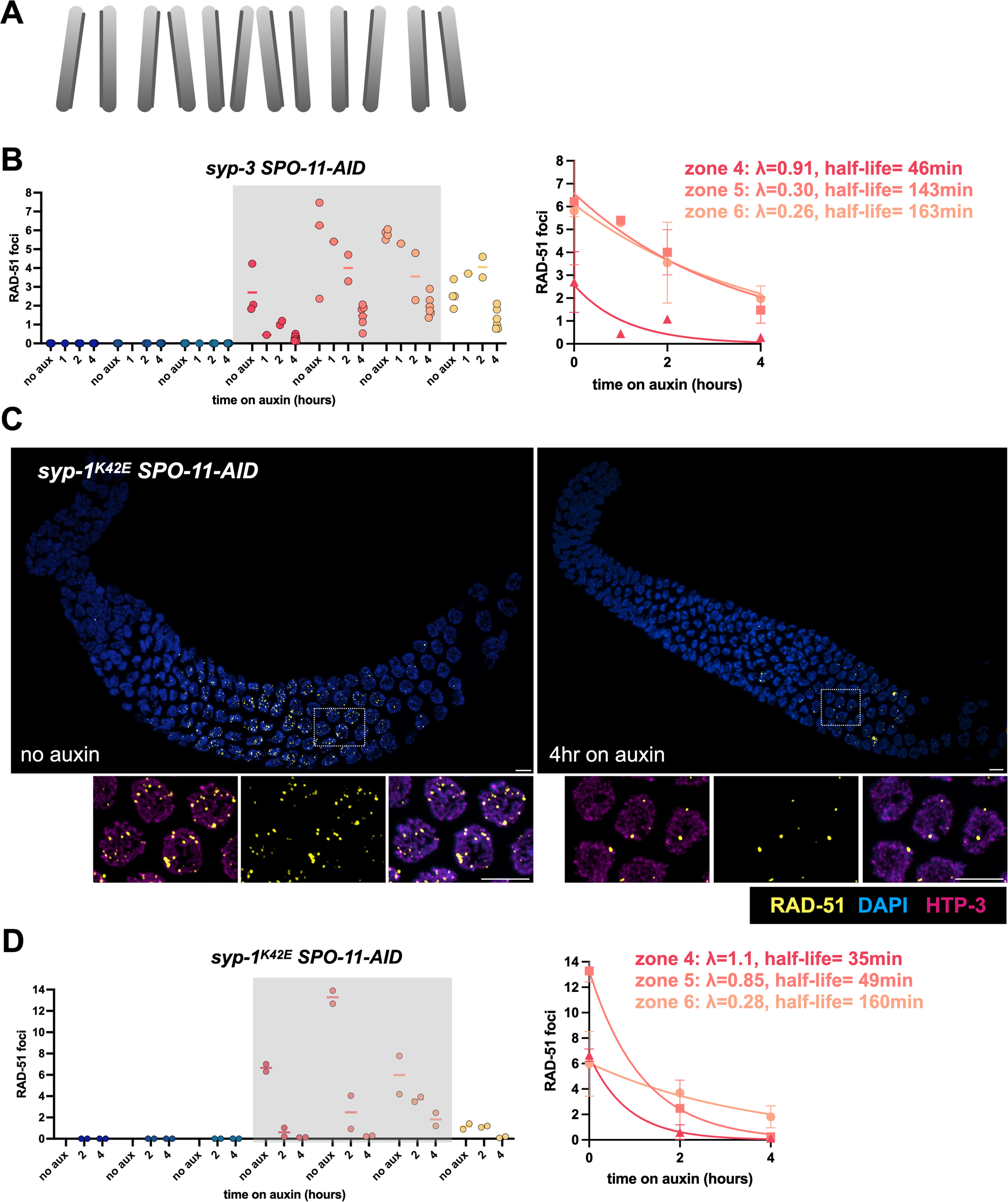
RAD-51 foci lifetime in *syp-3* and *syp-1^K42E^* worms. (A) Diagram of the karyotype in *syp-3* and *syp-1^K42E^* worms, where all chromosomes are not associated with their partner and cannot form crossovers. (B) Left, RAD-51 foci number per nucleus in *syp-3 SPO-11-AID* following 0, 1, 2 and 4 hours on auxin, colored by gonad zone as in Fig. 1. Grey shading indicates zones where decay rate was calculated. Right, exponential decay of RAD-51 foci number in zones 4, 5 and 6. Error bars indicate standard deviation. See Fig. S1 for immunofluorescence images. (C) Maximum intensity projection of whole gonads from *syp-1^K42E^ SPO-11-AID* worms stained for RAD-51 (yellow) and DNA (DAPI; blue) on auxin for 0 hr and 4 hr. Right, inset showing higher magnification of pachytene nuclei stained for RAD-51 (yellow), the axis component HTP-3 (magenta) and DNA (DAPI; blue). Scale bars = 5 μm. (D) Left, RAD-51 foci number per nucleus in *syp1^K42E^ SPO-11-AID* following 0, 2 and 4 hours on auxin, colored by gonad zone as in Fig. 1. Grey shading indicates zones where decay rate was calculated. Right, exponential decay of RAD-51 foci number in zones 4, 5 and 6. Error bars indicate standard deviation.

**Figure 4:**
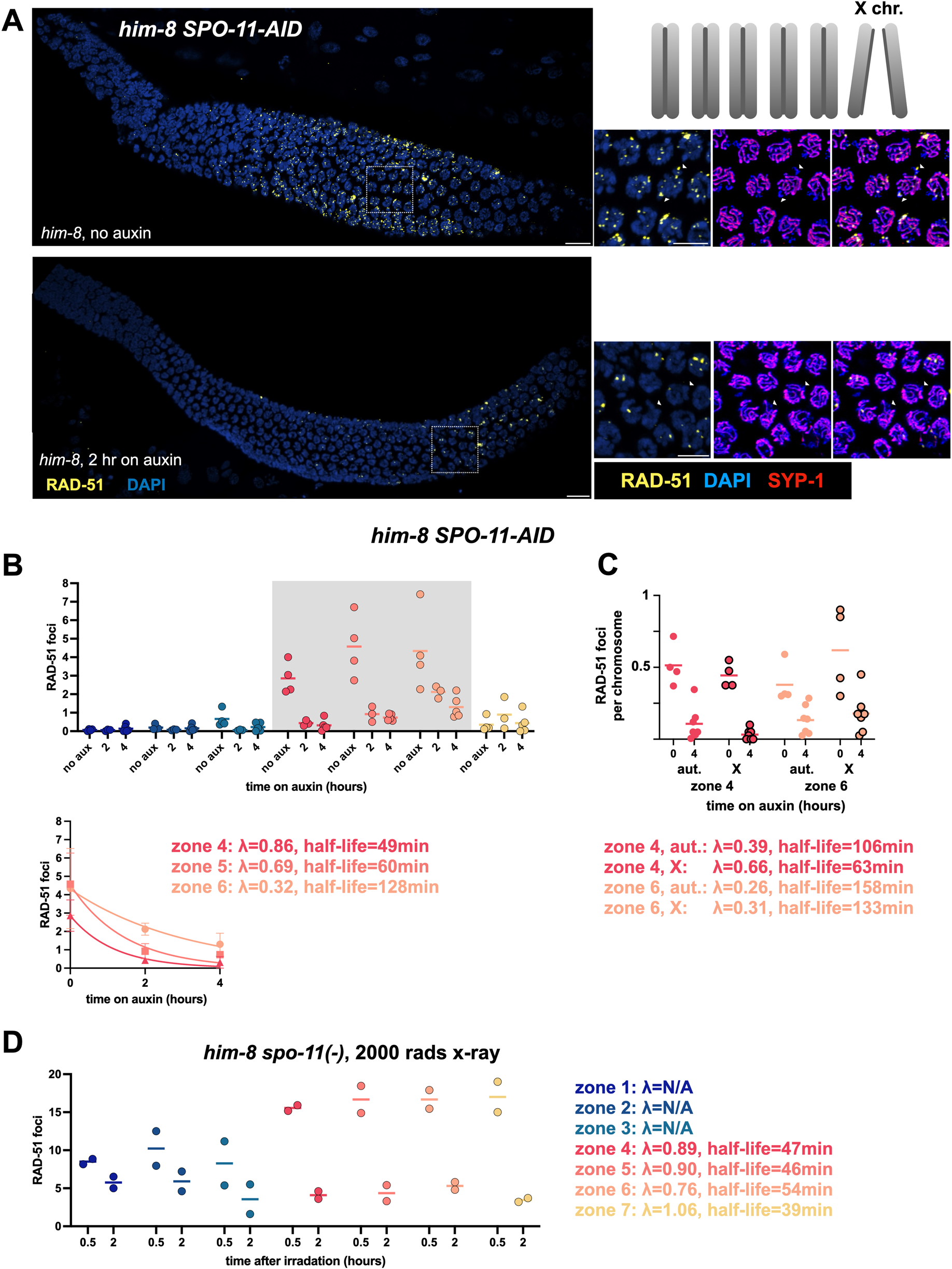
RAD-51 foci lifetime in *him-8* worms. (A) Maximum intensity projection of whole gonads from *him-8 SPO-11-AID* hermaphrodites stained with RAD-51 (yellow) and DAPI (blue). Right, inset showing higher magnification of pachytene nuclei stained with RAD-51 (yellow), the axis component HTP-3 (blue) and the synaptonemal complex component SYP-1 (red). The X chromosomes are identified as having axis staining but lacking synaptonemal complex staining (examples are shown with white triangles). Top right, diagram of the karyotype in *him-8* worms, where the X chromosome is not paired and cannot form crossovers. Scale bars = 5 μm. (B) Left, RAD-51 foci number per nucleus in *him-8 SPO-11-AID* animals following 0, 1, 2 and 4 hours on auxin, colored by gonad zone as in Fig. 1. Gray shading indicate zones where decay rate was calculated. Bottom, exponential decay of RAD-51 foci number in zones 4, 5 and 6. Error bars indicate standard deviation. (C) RAD-51 foci number per chromosome in *him-8 SPO-11-AID* animals following 0 and 4 hours on auxin in zones 4 and 6. The values for the autosomes (aut.) lack borders and the X chromosome noted by black borders. (D) RAD-51 foci number per nucleus in *him-8 spo-11(-)* animals 0.5 and 2 hours after irradiation, colored by gonad zone as in Fig. 1.

We also analyzed *syp-1^K42E^* worms, where synaptonemal complex material is present but it associates with unpaired chromosomes (24). (Since *syp-1^K42E^* is a temperature-sensitive mutation, this analysis was performed at 25°C). In *syp-1^K42E^* worms, RAD-51 foci exhibited similar kinetics upon SPO-11 shut-off: 35, 49, 160 minutes in zones 4, 5, and 6, respectively (Fig. 3C-D). The similar repair kinetics in control, *syp-3* and *syp-1^K42E^* worms suggests that strand-invasion proceeds at similar rates regardless of the repair template.

To ensure that the kinetics we measured in *syp-3* and *syp-1^K42E^* worms are not a consequence of the nucleus-wide failure to align the homologs, we analyzed conditions where homolog-directed repair is suppressed in only parts of the genome. We first analyzed strains where the associations between one pair of homologs are affected. In *him-8* mutant worms, the X chromosomes fail to pair and align (44). The starting number of RAD-51 foci is greater in *him-8* animals, and that they occur over a larger portion of the gonad, as previously reported (13, 44, 45). However, their disappearance following SPO-11 shut-off occurred at rates similar to controls (Fig. 4A-B). RAD-51 foci exhibited half-lives of 49, 60 and 128 mins in zones 4, 5 and 6, respectively. Furthermore, we found that RAD-51 foci on the unpaired X chromosomes *versus* the paired autosomes had similar half-lives (Fig. 4C). As an independent measure of strand-invasion kinetics we analyzed RAD-51 kinetics in irradiated *him-8 spo-11(-)* worms. We found similar RAD-51 foci kinetics compared with controls (Fig. 2 and 4), consistent with similar kinetics for homolog- and sister-directed repair upon SPO-11 shut-off.

We also attempted to analyze *zim-2* animals, where chromosome V is unable to pair (25). Like *him-8* worms, *zim-2* worms exhibit a higher overall number of RAD-51 foci (Fig. S2; (46)). However, in many nuclei RAD-51 persisted in 1-2 clusters, each larger than a diffraction-limited focus. These clusters were prevalent in later stages of meiosis, and mostly localized the unpaired chromosome V. (We occasionally observed similar clusters in *him-8* worms as well; see Fig. 4A.) The high prevalence of nuclei with these clusters, where individual foci could not be reproducibly counted, prevented quantitative analysis. Notably, we did not observe the clusters in irradiated *zim-2 spo-11(-)* worms, which exhibited RAD-51 kinetics indistinguishable from control and *him-8* worms (Fig. S2). We speculate that these clusters might reflect the coalescence of nearby repair intermediates. However, the reason for the higher prevalence of these clusters in *zim-2 versus him-8* worms is currently unclear (see Discussion).

Finally, we analyzed worms harboring a large inversion, *mIn1*. In *mIn1/+* heterozygous worms all chromosomes are aligned. However, approximately half of chromosome II is aligned with a non-homologous partner and can therefore only undergo sister-directed repair (see diagram in Fig. 5A). Similar to *him-8* worms, some nuclei harbored clustered RAD-51, as previously reported (20). After SPO-11 shut-off, we observed a rapid decrease in the number of RAD-51 foci, which exhibited half-lives of 64 and 138 in zones 4 and 5, respectively. Given that only sister-directed repair is possible in the inverted region in *mIn1/+* worms, decay kinetics in this strain suggests that sister- and homolog-directed strand-invasion occur with similar kinetics.

**Figure 5:**
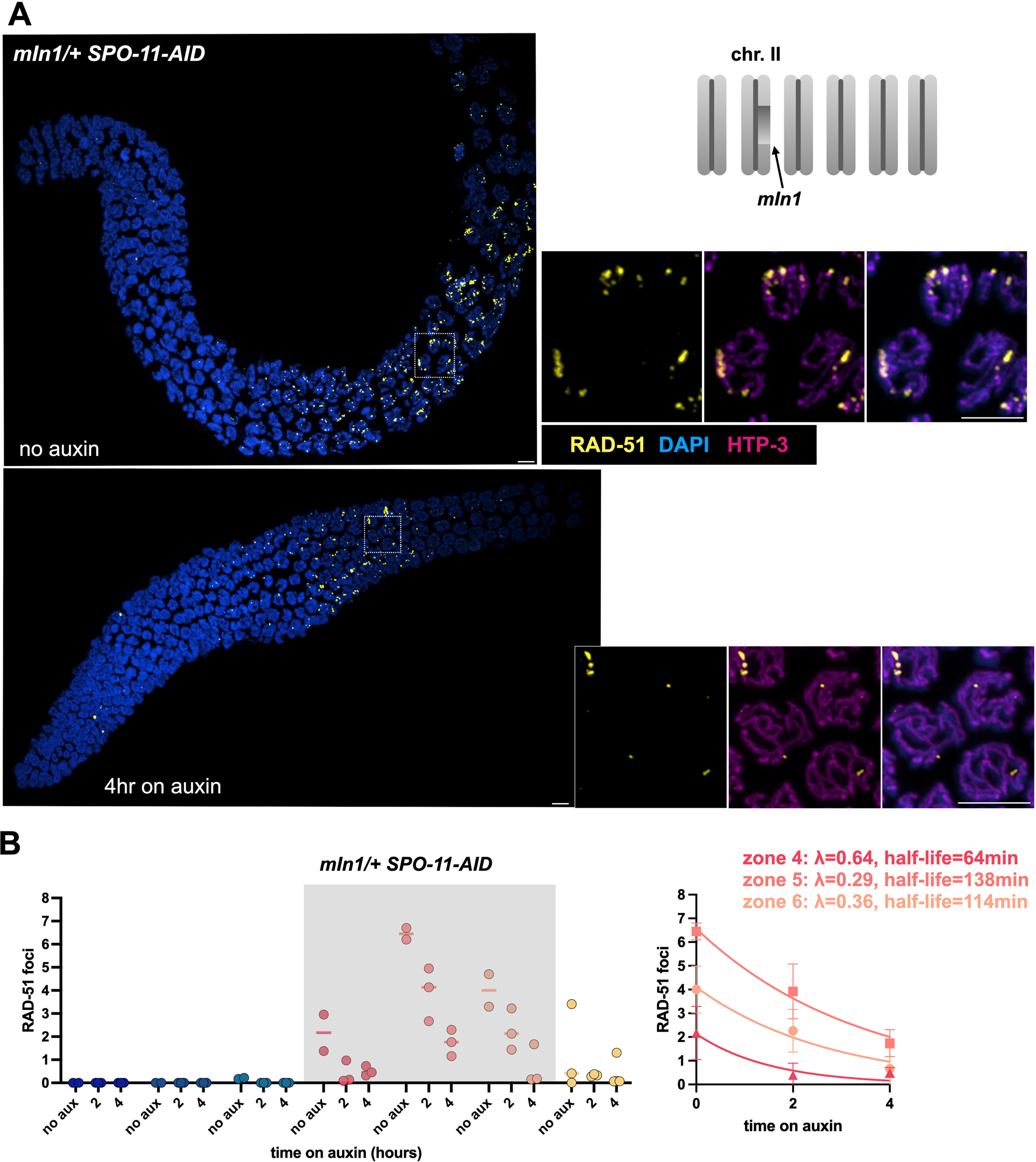
RAD-51 foci lifetime in *mIn1/+* worms. (A) Maximum intensity projection of whole gonads from *mIn1/+ SPO-11-AID* worms stained for RAD-51 (yellow) and DNA (DAPI; blue) on auxin for 0 and 4 hours. Right, inset showing higher magnification of pachytene nuclei, also showing staining for the axis component HTP-3 (magenta). Top right, diagram of the karyotype in *mIn1/+* worms, where all chromosomes are aligned but the inverted portion of chromosome II is paired non-homologously and cannot form crossovers. Scale bars = 5 μm. (B) Left, RAD-51 foci number per nucleus in *mIn1/+ SPO-11-AID* following 0, 2 and 4 hours on auxin, colored by gonad zone as in Fig. 1. Grey shading indicates zones where decay rate was calculated. Right, exponential decay of RAD-51 foci number in zones 4, 5 and 6. Error bars indicate standard deviation.

Taken together, our analysis indicates that RAD-51 foci lifetime - a proxy for repair kinetics - is similar in multiple conditions where some or all chromosomes are not paired and aligned. By extension, these observations suggest that the progression of sister-directed strand invasion is not inherently slowed down relative to homolog-directed repair.

### Calculating the total number of meiotic DSBs

The overall number of DSBs is crucial for our understanding of the mechanisms regulating meiotic DNA repair and crossover formation since it serves as the ’substrate’ for the various repair pathways. However, this number has proven elusive (47) despite great progress in our understanding of the structure and genetic dependencies of repair intermediates.

To derive the overall DSB number in meiosis, we used our calculated RAD-51 lifetimes. The change in RAD-51 foci number, *dC/dt*, is a result of the number of new DSBs per hour, *B*, and the number of intermediates that have undergone repair, *λC*, based on the number of RAD-51 intermediates, *C*, and the rate by which they are repaired (the exponential decay coefficient), *λ*:

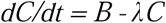

(More precisely, *B* is the number of processed DSBs that become associated with RAD-51 per hour; however, as discussed above, DSB processing time seems to be much shorter than one hour, and, therefore, *B* is well-approximated by the number of newly formed DSBs).

Within each zone, the average number of observed RAD-51 foci reflects the steady-state number of intermediates. In other words, *dC/dt = 0*, and consequently:

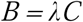

Based on this formula, the rate of DSB formation in zones 4 and 5 - where most DSBs occur - is 1.6 and 2.1 DSBs per hour, respectively. The time spent in each meiotic zone is 8.4, 8.3 and 6.9 hours for zones 4, 5 and 6, respectively. This estimate comes from the average movement of nuclei in the gonad (∼1 row/hour; (31–35, 48)) multiplied by the average number of rows in each zone. Together, these values yield a total of 40.1 DSBs per meiotic nucleus, or 6.7 DSBs per homolog pair (Fig. 6). This number is likely an underestimate. First, we only included DSBs occurring in zones 4 and 5, whereas some RAD-51 foci are also visible in zones 3, 6 and 7. Second, we likely overestimated RAD-51 lifetimes by not considering the time to degrade SPO-11 on auxin and to process the DSBs; shorter lifetimes will translate to more DSBs. Nonetheless, assuming 40 DSBs per nucleus, DSBs outnumber crossovers 7:1 in worms, in line with measurements in other organisms (4, 49).

**Figure 6:**
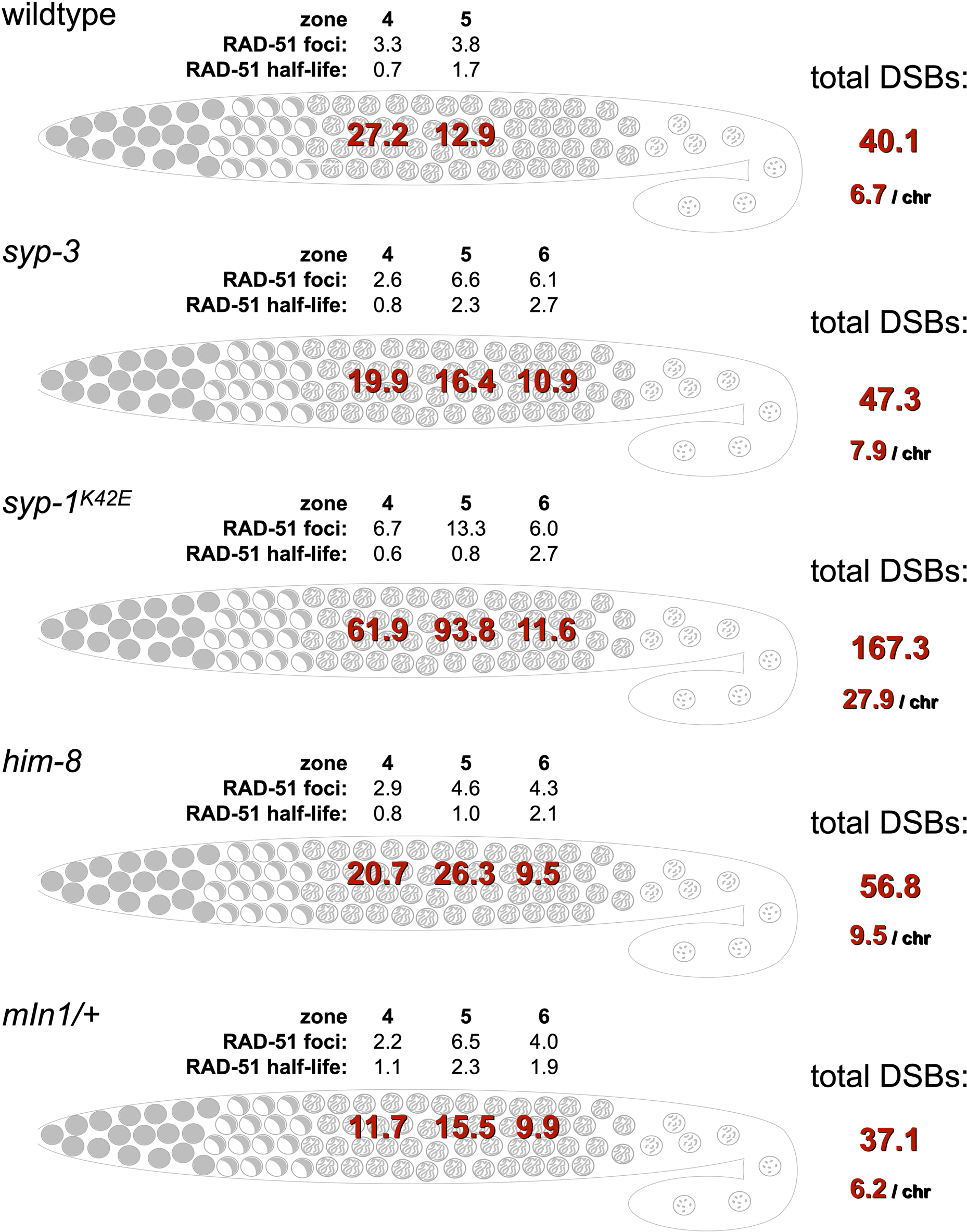
Overall DSB number based on RAD-51 number and strand-invasion kinetics. Calculated total DSBs per meiotic nuclei in wild-type, *syp-3, syp-1^K42E^*, *him-8,* and *mIn1/+* animals. RAD-51 foci numbers (based on the ’no auxin’ data) and half-lives (in hours) are taken from Figs. 1, 3, 4 and 5. See text for details.

The total number of DSBs is higher in most mutant scenarios that we tested. *syp-3* and *him-8* worms exhibit a higher number of RAD-51 foci, with each nucleus undergoing 46.5 and 56.8 DSBs, respectively. Nuclei in *mIn1/+* worms underwent similar number of DSBs to wild-type worms (37.1 DSBs). The presence of many RAD-51 foci on the unpaired or mis-paired chromosomes in fixed images of *him-8* and *mIn1/+* worms (Figs. 4 and 5), suggests that these chromosomes likely incur more DSBs throughout meiosis. This suggests that, for instance, the X chromosomes in *him-8* worms are likely to undergo upwards of 20 DSBs. *syp-1^K42E^* worms exhibited a remarkable 167 DSBs per nucleus, entailing almost 28 DSBs per chromosome. The exact reason for the large difference between the two conditions that do not align homologs across the genome - *syp-3* and *syp-1^K42E^* - is not clear. A possible explanation is the difference in meiotic progression, with nuclei in *syp-3* worms remaining in a leptotene-zygotene (the so-called ’transition zone’) and nuclei in *syp-1^K42E^* worms progressing to a pachytene-like state (24). A non-mutually exclusive possibility is that the increase in DSBs stems from activation of different checkpoints in the two mutants.

## Discussion

Here, we controlled DSB influx to analyze the kinetics of repair during *C. elegans* meiosis. We find that RAD-51-associated intermediates – which carry out the homology search and strand invasion stages of homologous recombination – have a half-life of 1-2 hours. This duration is mostly unaffected by meiotic stage, repair template, or DSB location.

One of the distinguishing features of meiotic DSB repair is the preferential use of the homolog as a repair template rather than the sister chromatid (4, 47). However, the extent of this “homolog bias”, and the molecular mechanism implementing it, remain poorly understood. In worms, unpaired and mis-paired chromosomes cause an overall increase in the number of RAD-51 foci, accumulation of foci on genomic regions where the homolog is inaccessible, and persistence of foci until later stages of meiosis (50–53). These observations raised the possibility that repair events that cannot engage the homolog ‘stall’ during much of meiotic prophase, until they are eventually channeled into sister-templated pathways, which serve as a backup repair mechanism (e.g., (19)). Such differential timing suggests that homolog bias might be implemented by arresting sister-engaged DSB repair intermediates while allowing homolog-directed intermediates to proceed. Our finding of similar repair kinetic for sister- and homolog-directed repair argue against that possibility. Instead, our work suggests that during meiosis unpaired chromosomes undergo multiple cycles of DSB formation and sister-directed repair.

While our work sheds light on the average kinetics of repair, we cannot rule out that a subset of events is regulated differently. In addition, we note that our calculations of the overall number of meiotic DSBs would be affected by difficult-to-control experimental conditions, such as antibody source, microscope settings and the thresholds used during image analysis, which could all yield variable number of foci. Such differences have indeed been observed (see (47) for further discussion). Importantly, our kinetic analysis, which detects foci in the same way at all time points, is mostly free of this potential source of error.

We and others have noted the formation of clusters of RAD-51 foci, particularly in genomic regions that cannot form crossovers (e.g., (13, 54)). Here, we have noted such clusters on the unpaired X chromosomes in *him-8* mutants (rarely; Fig. 4) and the unpaired chromosome V in *zim-2* mutants (commonly; Fig. S2). We also noted these cluster on the inverted regions in *mIn1/+* worms (Fig. 5). Several observations might provide hints as to the etiology of these clusters. First, clusters were not prominent in conditions where crossovers could not form genome-wide (*syp-3* and *syp-1^K42E^*; Fig. 3). Second, clusters were not observed upon irradiation in *zim-2* mutants that lack endogenous DSBs (Fig. S2). Third, clusters were more prominent in later stages of meiosis (Fig. S2). Finally, some of the clusters are still present at the 4 hours timepoints (Figs. 4 and 5). These observations are all consistent with RAD-51 clusters representing abnormal repair, perhaps as a result of a high local concentration of DSBs. Smaller number of DSBs on the X chromosome *versus* the autosomes could explain the higher prevalence of clusters in *zim-2* worms. The distribution of excess DSBs throughout the genome, rather than localized at a specific region, could explain why conditions such as *syp-3* and *syp-1^K42E^* do not form clusters as often. An important conclusion of our work is that these RAD-51 clusters are unlikely to stem from inherently slow DSB repair in the absence of the homolog. Nonetheless, the reason for the formation of RAD-51 clusters only in some genetic conditions, as well as the mechanisms that eventually resolves them, remain a subject of future investigation.

Our work has allowed us to quantify the overall number of DSBs by combining repair kinetics with the number of repair intermediates. This method joins other approaches that have been used to derive this number. One approach, which has been used extensively in yeast, follows the fate of a few heavily-used break sites to give population-level averages of different intermediates (55). However, the repair of DSBs genome-wide cannot be trivially extrapolated from a single DSB (56), and physical analysis of a single DSB has not been reported in multicellular organisms. In another approach, the total number of meiotic DSBs has been estimated by combining snapshots of RAD-51 foci with mutations that prevent repair progression (47, 57, 58), presumably trapping all repair intermediates. However, the failure to form crossovers and the persistence of repair intermediates activate meiotic checkpoints (59), likely feeding back to the DSB formation machinery (60). Interestingly, the number of DSBs calculated here - 40 DSBs per nucleus - is similar to the number of RAD-51 counted in mutants blocked downstream of strand-invasion (*rad-54*, reported in (47), as well as to the maximum number of foci observed for markers of intermediates that are thought to act downstream of RAD-51 (RPA; (20)).

The overall number of meiotic DSBs has important implications. First, DSB number is important for understanding the mechanism of repair, as it is the ‘substrate’ for most repair pathways. Second, while crossovers are regulated independently of DSBs – e.g., each worm chromosome undergoes exactly one crossover regardless of the number of DSBs (3) – DSBs themselves are also tightly regulated. Specifically, DSBs are downregulated by nearby DSBs through the action of the ATM and ATR kinases (51, 60–62); and by the formation of at least one crossover on every homolog pair (51, 60, 63). Our work suggests that in conditions where homolog associations are disrupted, the unpaired chromosomes undergo more DSBs than previously thought. Such upregulation of DSBs is a dangerous proposition since it increases the risk of improper repair – an important evolutionary cost of sexual reproduction (64).

## Acknowledgments

We would like to thank the Rog lab for discussions; Sarit Smolikove and Nicola Silva for sharing raw data; Kent Golic for discussions; and Yuval Mazor for comments on this manuscript. Some strains were provided by the Caenorhabditis Genetics Center, which is funded by NIH Office of Research Infrastructure Programs (P40 OD010440).

## Data Availability

Worm strains are available upon request. The authors affirm that all data necessary for confirming the conclusions of the article are present within the article and figures.

## References

1. Keeney, S., Giroux, C.N. and Kleckner, N. (1997) Meiosis-specific DNA double-strand breaks are catalyzed by Spo11, a member of a widely conserved protein family. Cell, 88, 375–384.

2. Dernburg, A.F., McDonald, K., Moulder, G., Barstead, R., Dresser, M. and Villeneuve, A.M. (1998) Meiotic recombination in C. elegans initiates by a conserved mechanism and is dispensable for homologous chromosome synapsis. Cell, 94, 387–398.

3. Yokoo, R., Zawadzki, K.A., Nabeshima, K., Drake, M., Arur, S. and Villeneuve, A.M. (2012) COSA-1 reveals robust homeostasis and separable licensing and reinforcement steps governing meiotic crossovers. Cell, 149, 75–87.

4. Zickler, D. and Kleckner, N. (2023) Meiosis: Dances Between Homologs. Annu. Rev. Genet., 10.1146/annurev-genet-061323-044915.

5. Clejan, I., Boerckel, J. and Ahmed, S. (2006) Developmental modulation of nonhomologous end joining in Caenorhabditis elegans. Genetics, 173, 1301–1317.

6. 6. Adamo, A., Collis, S.J., Adelman, C.A., Silva, N., Horejsi, Z., Ward, J.D., Martinez-Perez, E., Boulton, S.J. and La Volpe, A. (2010) Preventing nonhomologous end joining suppresses DNA repair defects of Fanconi anemia. Mol. Cell, 39, 25–35.

7. Smolikov, S., Eizinger, A., Hurlburt, A., Rogers, E., Villeneuve, A.M. and Colaiácovo, M.P. (2007) Synapsis-defective mutants reveal a correlation between chromosome conformation and the mode of double-strand break repair during Caenorhabditis elegans meiosis. Genetics, 176, 2027–2033.

8. Martin, J.S., Winkelmann, N., Petalcorin, M.I.R., McIlwraith, M.J. and Boulton, S.J. (2005) RAD-51-dependent and -independent roles of a Caenorhabditis elegans BRCA2-related protein during DNA double-strand break repair. Mol. Cell. Biol., 25, 3127–3139.

9. Almanzar, D.E., Gordon, S.G. and Rog, O. (2021) Meiotic sister chromatid exchanges are rare in C. elegans. Curr. Biol., 31, 1499–1507.e3.

10. Toraason, E., Horacek, A., Clark, C., Glover, M.L., Adler, V.L., Premkumar, T., Salagean, A., Cole, F. and Libuda, D.E. (2021) Meiotic DNA break repair can utilize homolog-independent chromatid templates in C. elegans. Curr. Biol., 31, 1508–1514.e5.

11. Jackson, J.A. and Fink, G.R. (1985) MEIOTIC RECOMBINATION BETWEEN DUPLICATED GENETIC ELEMENTS IN *SACCHAROMYCES CEREVISIAE*. Genetics, 109, 303–332.

12. Schwacha, A. and Kleckner, N. (1997) Interhomolog bias during meiotic recombination: Meiotic functions promote a highly differentiated interhomolog-only pathway. Cell, 90, 1123–1135.

13. MacQueen, A.J., Phillips, C.M., Bhalla, N., Weiser, P., Villeneuve, A.M. and Dernburg, A.F. (2005) Chromosome sites play dual roles to establish homologous synapsis during meiosis in C. elegans. Cell, 123, 1037–1050.

14. Goldfarb, T. and Lichten, M. (2010) Frequent and efficient use of the sister chromatid for DNA double-strand break repair during budding yeast meiosis. PLoS Biol., 8, e1000520.

15. Haber, J.E., Thorburn, P.C. and Rogers, D. (1984) Meiotic and mitotic behavior of dicentric chromosomes in Saccharomyces cerevisiae. Genetics, 106, 185–205.

16. Borde, V. (2007) The multiple roles of the Mre11 complex for meiotic recombination. Chromosome Res., 15, 551–563.

17. 17. Rinaldo, C., Bazzicalupo, P., Ederle, S., Hilliard, M. and La Volpe, A. (2002) Roles for Caenorhabditis elegans rad-51 in meiosis and in resistance to ionizing radiation during development. Genetics, 160, 471–479.

18. Alpi, A., Pasierbek, P., Gartner, A. and Loidl, J. (2003) Genetic and cytological characterization of the recombination protein RAD-51 in Caenorhabditis elegans. Chromosoma, 112, 6–16.

19. 19. Colaiácovo, M.P., MacQueen, A.J., Martinez-Perez, E., McDonald, K., Adamo, A., La Volpe, A. and Villeneuve, A.M. (2003) Synaptonemal complex assembly in C. elegans is dispensable for loading strand-exchange proteins but critical for proper completion of recombination. Dev. Cell, 5, 463–474.

20. Woglar, A. and Villeneuve, A.M. (2018) Dynamic architecture of DNA repair complexes and the synaptonemal complex at sites of meiotic recombination. Cell, 173, 1678–1691.e16.

21. Hayashi, M., Chin, G.M. and Villeneuve, A.M. (2007) C. elegans germ cells switch between distinct modes of double-strand break repair during meiotic prophase progression. PLoS Genet., 3, e191.

22. Brenner, S. (1974) The genetics of Caenorhabditis elegans. Genetics, 77, 71–94.

23. Zhang, L., Ward, J.D., Cheng, Z. and Dernburg, A.F. (2015) The auxin-inducible degradation (AID) system enables versatile conditional protein depletion in C. elegans. Development, 142, 4374–4384.

24. Gordon, S.G., Kursel, L.E., Xu, K. and Rog, O. (2021) Synaptonemal Complex dimerization regulates chromosome alignment and crossover patterning in meiosis. PLoS Genet., 17, e1009205.

25. Phillips, C.M. and Dernburg, A.F. (2006) A family of zinc-finger proteins is required for chromosome-specific pairing and synapsis during meiosis in C. elegans. Dev. Cell, 11, 817–829.

26. Phillips, C.M., McDonald, K.L. and Dernburg, A.F. (2009) Cytological analysis of meiosis in Caenorhabditis elegans. Methods Mol. Biol., 558, 171–195.

27. Hurlock, M.E., Čavka, I., Kursel, L.E., Haversat, J., Wooten, M., Nizami, Z., Turniansky, R., Hoess, P., Ries, J., Gall, J.G., et al. (2020) Identification of novel synaptonemal complex components in *C. elegans*. J. Cell Biol., 219.

28. Harper, N.C., Rillo, R., Jover-Gil, S., Assaf, Z.J., Bhalla, N. and Dernburg, A.F. (2011) Pairing centers recruit a polo-like kinase to orchestrate meiotic chromosome dynamics in C. elegans. Dev. Cell, 21, 934–947.

29. Hicks, T., Trivedi, S., Eppert, M., Bowman, R., Tian, H., Dafalla, A., Crahan, C., Smolikove, S. and Silva, N. (2022) Continuous double-strand break induction and their differential processing sustain chiasma formation during Caenorhabditis elegans meiosis. Cell Rep., 40, 111403.

30. Zhang, L., Köhler, S., Rillo-Bohn, R. and Dernburg, A.F. (2018) A compartmentalized signaling network mediates crossover control in meiosis. Elife, 7.

31. Hubbard, E.J.A. and Schedl, T. (2019) Biology of the Caenorhabditis elegans Germline Stem Cell System. Genetics, 213, 1145–1188.

32. Fox, P.M. and Schedl, T. (2015) Analysis of Germline Stem Cell Differentiation Following Loss of GLP-1 Notch Activity in Caenorhabditis elegans. Genetics, 201, 167–184.

33. Mlynarczyk-Evans, S. and Villeneuve, A.M. (2017) Time-Course Analysis of Early Meiotic Prophase Events Informs Mechanisms of Homolog Pairing and Synapsis in Caenorhabditis elegans. Genetics, 207, 103–114.

34. Jaramillo-Lambert, A., Ellefson, M., Villeneuve, A.M. and Engebrecht, J. (2007) Differential timing of S phases, X chromosome replication, and meiotic prophase in the C. elegans germ line. Dev. Biol., 308, 206–221.

35. Crittenden, S.L., Leonhard, K.A., Byrd, D.T. and Kimble, J. (2006) Cellular analyses of the mitotic region in the Caenorhabditis elegans adult germ line. Mol. Biol. Cell, 17, 3051–3061.

36. Hunter, N. and Kleckner, N. (2001) The single-end invasion: an asymmetric intermediate at the double-strand break to double-holliday junction transition of meiotic recombination. Cell, 106, 59–70.

37. Ahuja, J.S., Sandhu, R., Huang, L., Klein, F. and Valentin Börner, G. (2024) Temporal and Functional Relationship between Synaptonemal Complex Morphogenesis and Recombination during Meiosis. bioRxiv, 10.1101/2024.01.11.575218.

38. Lemmens, B.B.L.G. and Tijsterman, M. (2011) DNA double-strand break repair in Caenorhabditis elegans. Chromosoma, 120, 1–21.

39. Prieler, S., Penkner, A., Borde, V. and Klein, F. (2005) The control of Spo11’s interaction with meiotic recombination hotspots. Genes Dev., 19, 255–269.

40. Keeney, S. (2008) Spo11 and the formation of DNA double-strand breaks in meiosis. Genome Dyn. Stab., 2, 81–123.

41. Rog, O. and Dernburg, A.F. (2013) Chromosome pairing and synapsis during Caenorhabditis elegans meiosis. Curr. Opin. Cell Biol., 25, 349–356.

42. Page, S.L. and Hawley, R.S. (2004) The genetics and molecular biology of the synaptonemal complex. Annu. Rev. Cell Dev. Biol., 20, 525–558.

43. MacQueen, A.J., Colaiácovo, M.P., McDonald, K. and Villeneuve, A.M. (2002) Synapsis-dependent and -independent mechanisms stabilize homolog pairing during meiotic prophase in C. elegans. Genes Dev., 16, 2428–2442.

44. Phillips, C.M., Wong, C., Bhalla, N., Carlton, P.M., Weiser, P., Meneely, P.M. and Dernburg, A.F. (2005) HIM-8 binds to the X chromosome pairing center and mediates chromosome-specific meiotic synapsis. Cell, 123, 1051–1063.

45. Carlton, P.M., Farruggio, A.P. and Dernburg, A.F. (2006) A link between meiotic prophase progression and crossover control. PLoS Genet., 2, e12.

46. Almanzar, D.E., Gordon, S.G., Bristow, C., Hamrick, A., von Diezmann, L., Liu, H. and Rog, O. (2023) Meiotic DNA exchanges in C. elegans are promoted by proximity to the synaptonemal complex. Life Sci Alliance, 6.

47. Rosu, S., Libuda, D.E. and Villeneuve, A.M. (2011) Robust crossover assurance and regulated interhomolog access maintain meiotic crossover number. Science, 334, 1286–1289.

48. Almanzar, D.E., Hamrick, A. and Rog, O. (2022) Single-sister labeling in the C. elegans germline using the nucleotide analog EdU. STAR Protocols, 3, 101344.

49. 49. de Massy, B. (2013) Initiation of meiotic recombination: how and where? Conservation and specificities among eukaryotes. Annu. Rev. Genet., 47, 563–599.

50. Baudrimont, A., Penkner, A., Woglar, A., Machacek, T., Wegrostek, C., Gloggnitzer, J., Fridkin, A., Klein, F., Gruenbaum, Y., Pasierbek, P., et al. (2010) Leptotene/zygotene chromosome movement via the SUN/KASH protein bridge in Caenorhabditis elegans. PLoS Genet., 6, e1001219.

51. Stamper, E.L., Rodenbusch, S.E., Rosu, S., Ahringer, J., Villeneuve, A.M. and Dernburg, A.F. (2013) Identification of DSB-1, a protein required for initiation of meiotic recombination in Caenorhabditis elegans, illuminates a crossover assurance checkpoint. PLoS Genet., 9, e1003679.

52. Zalevsky, J., MacQueen, A.J., Duffy, J.B., Kemphues, K.J. and Villeneuve, A.M. (1999) Crossing over during Caenorhabditis elegans meiosis requires a conserved MutS-based pathway that is partially dispensable in budding yeast. Genetics, 153, 1271–1283.

53. Kelly, K.O., Dernburg, A.F., Stanfield, G.M. and Villeneuve, A.M. (2000) Caenorhabditis elegans msh-5 is required for both normal and radiation-induced meiotic crossing over but not for completion of meiosis. Genetics, 156, 617–630.

54. Woglar, A., Yamaya, K., Roelens, B., Boettiger, A., Köhler, S. and Villeneuve, A.M. (2020) Quantitative cytogenetics reveals molecular stoichiometry and longitudinal organization of meiotic chromosome axes and loops. PLoS Biol., 18, e3000817.

55. Schwacha, A. and Kleckner, N. (1994) Identification of joint molecules that form frequently between homologs but rarely between sister chromatids during yeast meiosis. Cell, 76, 51–63.

56. Medhi, D., Goldman, A.S. and Lichten, M. (2016) Local chromosome context is a major determinant of crossover pathway biochemistry during budding yeast meiosis. Elife, 5.

57. Mets, D.G. and Meyer, B.J. (2009) Condensins regulate meiotic DNA break distribution, thus crossover frequency, by controlling chromosome structure. Cell, 139, 73–86.

58. Nottke, A.C., Beese-Sims, S.E., Pantalena, L.F., Reinke, V., Shi, Y. and Colaiácovo, M.P. (2011) SPR-5 is a histone H3K4 demethylase with a role in meiotic double-strand break repair. Proceedings of the National Academy of Sciences, 108, 12805–12810.

59. Yu, Z., Kim, Y. and Dernburg, A.F. (2016) Meiotic recombination and the crossover assurance checkpoint in Caenorhabditis elegans. Semin. Cell Dev. Biol., 54, 106–116.

60. Rosu, S., Zawadzki, K.A., Stamper, E.L., Libuda, D.E., Reese, A.L., Dernburg, A.F. and Villeneuve, A.M. (2013) The C. elegans DSB-2 protein reveals a regulatory network that controls competence for meiotic DSB formation and promotes crossover assurance. PLoS Genet., 9, e1003674.

61. Guo, H., Stamper, E.L., Sato-Carlton, A., Shimazoe, M.A., Li, X., Zhang, L., Stevens, L., Tam, K.C.J., Dernburg, A.F. and Carlton, P.M. (2022) Phosphoregulation of DSB-1 mediates control of meiotic double-strand break activity. Elife, 11.

62. Garcia, V., Gray, S., Allison, R.M., Cooper, T.J. and Neale, M.J. (2015) Tel1ATM-mediated interference suppresses clustered meiotic double-strand-break formation. Nature, 520, 114–118.

63. Mu, X., Murakami, H., Mohibullah, N. and Keeney, S. (2020) Chromosome-autonomous feedback down-regulates meiotic DNA break competence upon synaptonemal complex formation. Genes Dev., 34, 1605–1618.

64. Otto, S.P. and Payseur, B.A. (2019) Crossover Interference: Shedding Light on the Evolution of Recombination. Annu. Rev. Genet., 53, 19–44.

