## Supplemental Figures for "Kinetic analysis of strand invasion during *C. elegans* meiosis reveals similar rates of sister- and homolog-directed repair"

### Figure S1

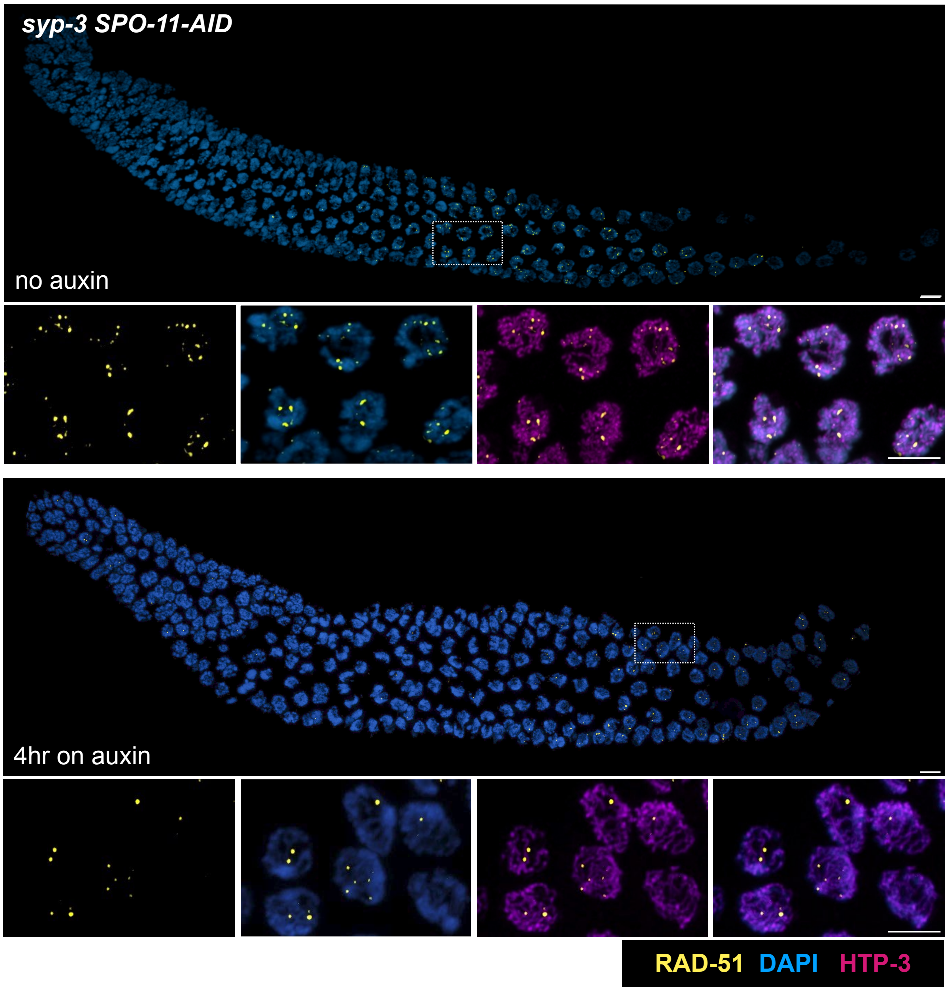


**Figure S1: RAD-51 staining in *syp-3 SPO-11-AID* worms**

Maximum intensity projection of whole gonads from *syp-3 SPO-11-AID* worms stained for RAD-51 (yellow) and DNA (DAPI; blue) on auxin for 0 hr and 4 hr. Right, inset showing higher magnification of pachytene nuclei stained for RAD-51 (yellow), the axis component HTP-3 (magenta), and DNA (DAPI; blue). Scale bars = 5 µm.

### Figure S2

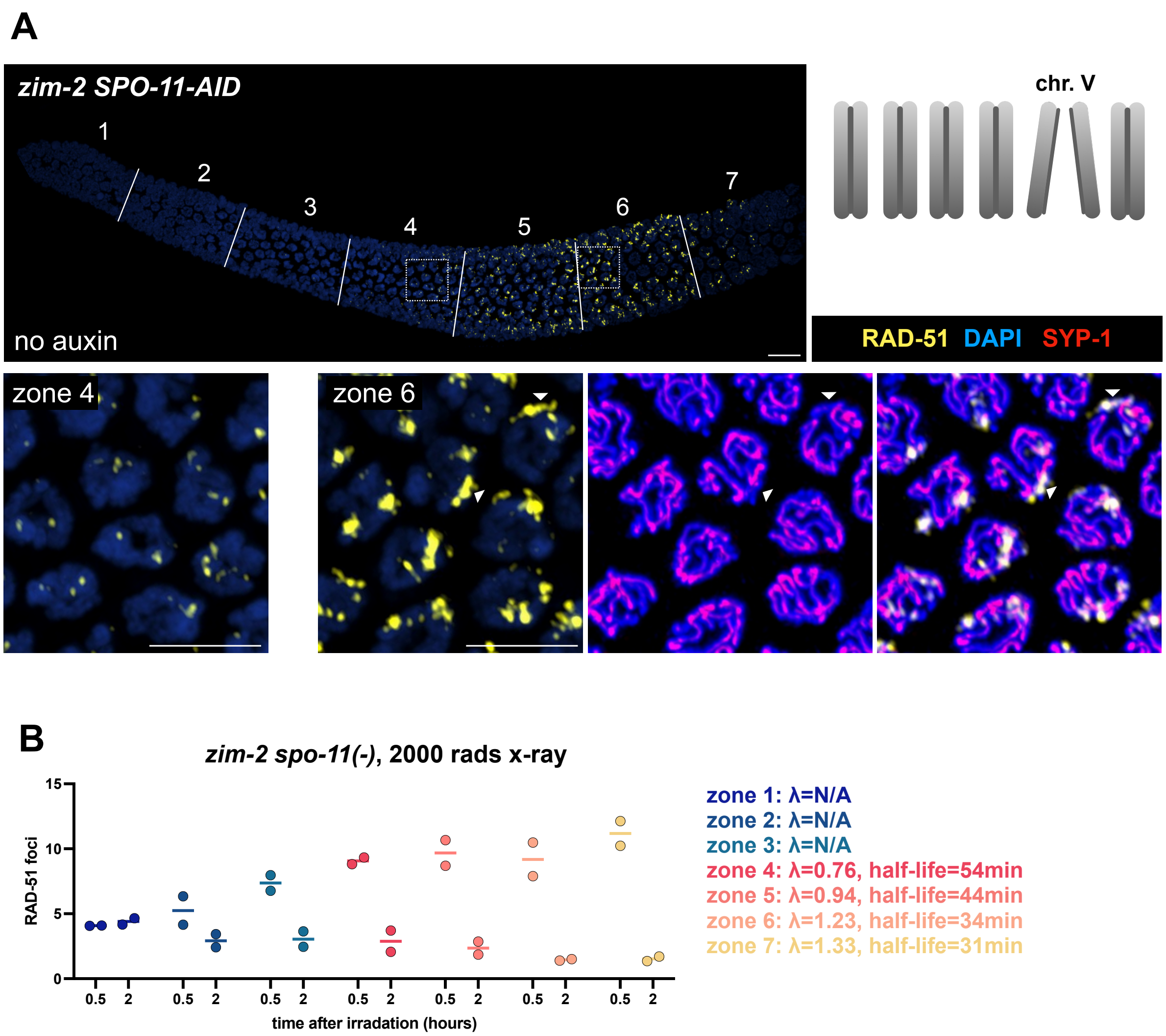


**Figure S2: RAD-51 foci analysis in *zim-2* worms**

(A) Maximum intensity projection of whole gonads from *zim-2 SPO-11-AID* hermaphrodites stained with RAD-51 (yellow) and DAPI (blue). Bottom left, an inset showing higher magnification of pachytene nuclei from zone 4. Bottom right, an inset showing higher magnification of pachytene nuclei from zone 6, stained with RAD-51 (yellow), the axis component HTP-3 (blue) and the synaptonemal complex component SYP-1 (red). Many of the large RAD-51 patches present in late pachytene localize to the unpaired chromosome V, identified as having axis staining but lacking synaptonemal complex staining (examples are shown with white triangles). Top right, diagram of the karyotype in *zim-2* worms, where chromosome V is not paired and cannot form crossovers. Scale bars = 5 µm. (B) RAD-51 foci number per nucleus in *zim-2 spo-11(-)* animals 0.5 and 2 hours after irradiation, colored by gonad zone as in Fig. 1.
